# Multidimensional behavioral profiles associated with resilience and susceptibility after inescapable stress

**DOI:** 10.1101/2023.11.08.566266

**Authors:** Benedito Alves de Oliveira-Júnior, Danilo Benette Marques, Matheus Teixeira Rossignoli, Tamiris Prizon, João Pereira Leite, Rafael Naime Ruggiero

## Abstract

Stress-related neuropsychiatric disorders are complex conditions that are difficult to diagnose and treat due to the variety of symptoms they present with. While animal models have been instrumental in understanding their underlying mechanisms, there is an ongoing debate about the validity of behavioral measures to address specific disorders. Here, we conducted an extensive ethological characterization of behavioral variables associated with anxiety and depression and employed a multidimensional approach to investigate stress susceptibility in male Wistar rats. The rats underwent inescapable footshocks (IS) or no-shocks (NS), followed by a comprehensive battery of behavioral tests over several days. Our study identified phenotypically distinct clusters, including fully characterized stress-susceptible and stress-resilient subpopulations, along with intermediate phenotypes. Reduced time in elevated plus maze open arms, diminished sucrose preference and increased escape failures consistently differentiated susceptible from resilient rats. Behavior in the forced swim test was unrelated to stress susceptibility. In conclusion, our study sheds light on the relationships between classical behavioral measures in animal models of depression, highlighting distinct patterns that differentiate susceptibility from resilience. It emphasizes the importance of a multidimensional assessment of behavioral responses in animal models for a better understanding of stress-related neuropsychiatric disorders.

**HIGHLIGHTS:**

- Clustering of features reveals a correlation pattern that separates behavioral variables associated with susceptibility from those associated with resilience to stress
- Multidimensional patterns of covariation among behavioral measures distinguish helpless from non-helpless individuals and indicate independence from forced swim test behavior.
- Clustering of individuals discriminates stressed subjects and reveals multidimensional behavioral phenotypes of resilience and susceptibility, indicating different stress-coping strategies.

## Introduction

Depressive disorders, as one of the main stress-related neuropsychiatric disorders, are the leading cause of disability worldwide, affecting more than 320 million people (WHO, 2017) and imposing a substantial economic burden (Greenberg, 2015). These disorders are complex and symptomatically heterogeneous conditions whose treatment is still ineffective in several clinical subtypes, many of which are unknown (Blackburn, 2019). Understanding the neurobiology underlying these conditions is crucial for developing more effective therapeutic interventions. In this context, modeling depressive symptoms in animal models has been recognized as an essential approach (Nestler & Hyman, 2010).

The majority of studies investigating depressive aspects in animal models focus on stress-induced models (Charney & Manji, 2004; Alfonso et al., 2005; Leigh, 2008). These models typically employ well-established experimental paradigms involving physical stressors (e.g., foot shocks) or psychosocial stressors (e.g., social defeat), followed by behavioral tests to assess alterations in depressive-related behaviors. Classical behavioral tests, such as the forced swim test and sucrose preference test, are commonly employed in animal models of depression due to their established predictive validity. However, this validity may not be related to the neurobiology of depression (Commons et al., 2017). In this regard, it is questioned whether some of these tests exclusively measure behaviors associated with depressive alterations. (Molendijk & De Kloet, 2015).

In the literature, even behavioral changes traditionally associated with a depressive-like profile due to stress are not consistent. Stepanichev et al. (2016) observed a decrease in sucrose consumption and preference, which was not correlated with immobility in the forced swim test. Additionally, Naudon & Jay (2015) identified distinct associations between immobility in the forced swim test and cognitive performance, as well as anxiety-related behaviors. In this sense, the punctual use of behavioral tests to distinguish the depressive phenotype seems to generate inconsistent results. In contrast, studying different behavioral categories associated with depressive-like behaviors is essential to understanding the underlying neural patterns (Marques et al., 2022) and gaining a better comprehension of the modeled phenomenon. For this purpose, a multivariate approach to behavioral assessment, using a set of behavioral tests, can be a valuable alternative. This methodology has shown promise in understanding the relations between behavioral domains in naive animals (Ramos et al., 1997; Feyissa et al., 2017) and in animal models of stress-related neuropsychiatric disorders (Kanari et al., 2005; Saénz et al., 2006), enriching the ethological assessment of the models, as well as their translational aspects. Despite this, there are still few studies that use this approach.

This study aimed to understand the association between a wide range of behavioral measures and investigate how their covariations characterize the depressive-like phenotype within the population. Furthermore, we aimed to examine the predictive value of these behavioral measures in determining susceptibility or resilience to stress in rats exposed to acute inescapable footshocks (IS) compared to those exposed to no shocks (NS). We found that behavioral variables exhibit a correlation pattern that distinguishes variables related to susceptibility or resilience to stress. Furthermore, we identified a covariation pattern consisting of decreased time spent in the open arms of the elevated plus maze, reduced sucrose preference, and elevated escape failures, consistently distinguishing helpless from not helpless rats. Notably, the behavior observed in the forced swim test did not exhibit a significant correlation with these behavioral profiles. Additionally, we discerned distinct phenotypic clusters, including subpopulations entirely characterized by susceptibility or resilience to stress, thus indicating varying stress-coping strategies.

## Results

### A single-day session of acute inescapable shocks promotes long-term helplessness and discriminates between helpless (H) and not helpless (NH) individuals

We performed a multidimensional investigation of behavioral measures traditionally related to depression and anxiety throughout a test battery after a single session of acute stress (Fig. 1). First, we assessed the univariate results by comparing the IS and NS groups. Figure 2 presents the outcomes of the main behavioral measures indicating that the IS group exhibited lower escape performance in the shuttle box than the NS group (escape failures: U = 693.5, p < 0.0001; latency to escape: U = 743, p = 0.0013; Fig. 2A). Additionally, there is a significant difference between the groups in the acoustic startle test (U = 627, p = 0.0342). Interestingly, no differences between groups were found in other behavioral measures traditionally related to depressive-like symptoms, such as immobility time in the forced swim test (t_(43)_ = −0.0993, p = 0.9213), social interaction (t_(43)_ = 0.0618, p = 0.9509), or sucrose preference (t_(43)_ = −1.5454, p = 0.1295).

**Figure 1.**
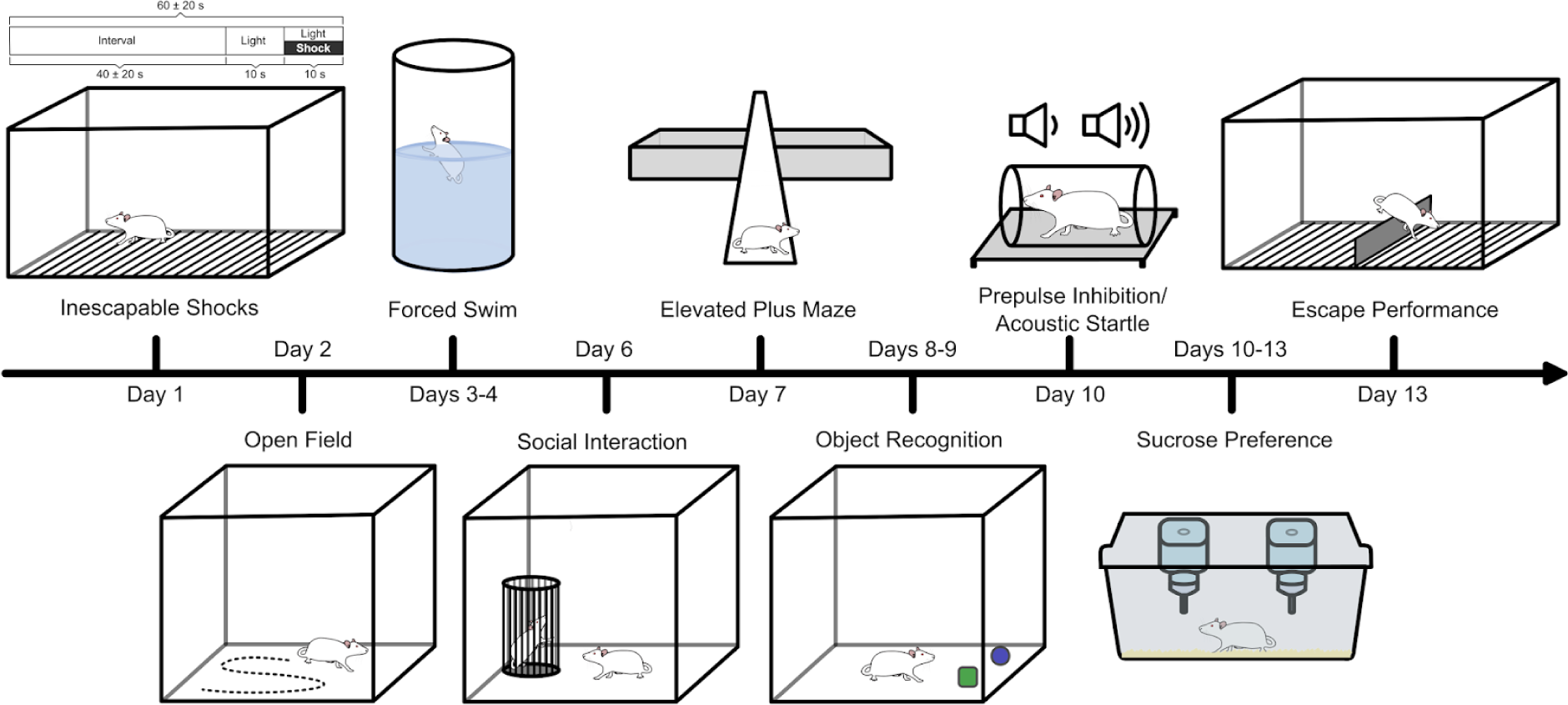
Behavioral tests. Schematic illustration of the procedure for exposure to inescapable footshock and behavioral testing. Illustration made with inkscape software package (Inkscape Project, 2020).

**Figure 2.**
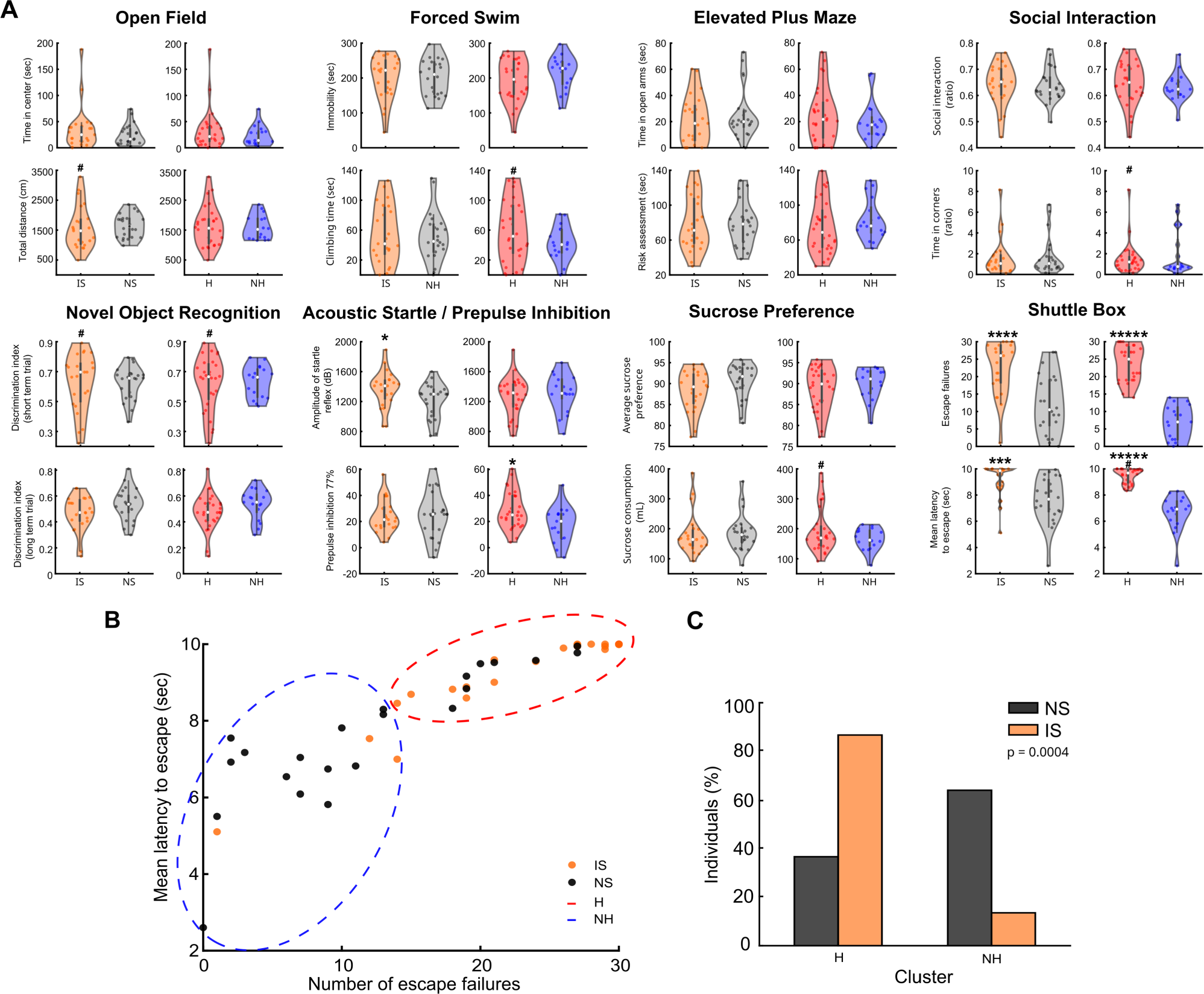
A single-day session of acute inescapable shocks promotes long-term helplessness and discriminates between helpless (H) and not helpless (NH) individuals. (**A**) Representative behavioral measures of each test. Comparison between groups IS (n=23) vs. NS (n=22), as well as H (n=28) vs. NH (n=17) clusters. (**B**) Unsupervised k-means clustering discriminates between H and NH clusters. (**C**) Individuals composing H and NH clusters. Student t-test: *p<0.05, ***p<0.001. Levene’s test: #p<0.05. Data in box plot shown as median +-1.5 × Interquatile.

Both groups exhibited wide variability in behavioral responses. During the evaluation of escape performance in the shuttle box, individuals in each group presented instances of both low escape performance, which is indicative of helplessness, and high escape performance, which is indicative of not being helpless. This variability suggests a predisposition of individuals to be susceptible or resilient to stress. To evaluate behavioral changes associated with stress predisposition, we employed escape performance in the shuttle box to classify individuals in the general population as either ‘H’ (helpless) or ‘NH’ (non-helpless). To achieve this, we employed an unsupervised clustering analysis using the k-means algorithm, where the optimal number of clusters was determined as two clusters (Fig. 2B). The analysis revealed that the H cluster encompassed 51.1% of the population (n = 23/45, X^2^_(1)_ = 12.2443, p = 0.0004, chi-squared test, Fig. 2C), with the majority of these individuals (86.9%, n = 20/23) coming from the IS group. Notably, the NH cluster accounted for 48.9% of the population (n = 22/45) and primarily consisted of NS individuals (63.6%, n = 14/22). Accordingly, NS individuals in the H cluster represent 13.1% (n = 3/23) and IS individuals in NS represent 36.3% (n = 8/22).

### Feature clustering distinguishes behavioral variables associated with susceptibility and resilience to stress

In order to evaluate how the most relevant variables from all behavioral tests are related, we measured the linear dependency between them using Spearman’s correlation and performed hierarchical clustering based on Euclidean Distance. The dendrogram generated by hierarchical clustering indicates the formation of two major clusters (Cophenetic coefficient: 0.6182, Fig. 3A), where variables whose increase is associated with stress resilience, such as climbing in forced swim and time in open arms of EPM, are grouped differently from those whose increase is associated with stress susceptibility, such as immobility in forced swim and time in closed arms of EPM (Fig. 3B). This result reveals two major covariance patterns of the data that significantly consisted of susceptibility- or resilience-related variables (X^2^_(1)_ = 15.0818, p = 0.0001, Fig. 3C). Notably, through a data permutation approach, where 70% of the data was randomly selected, it was confirmed that the separation between susceptibility- and resilience-related variables occurs significantly in most of the iterations with partitioned data (permutation of observations: 62.96% significant; permutation of variables: 62.25% significant, Fig. 3D) but not with randomized data (permutation of observations: 0.15% significant; permutation of variables: 0.46% significant, Fig. 3D).

**Figure 3.**
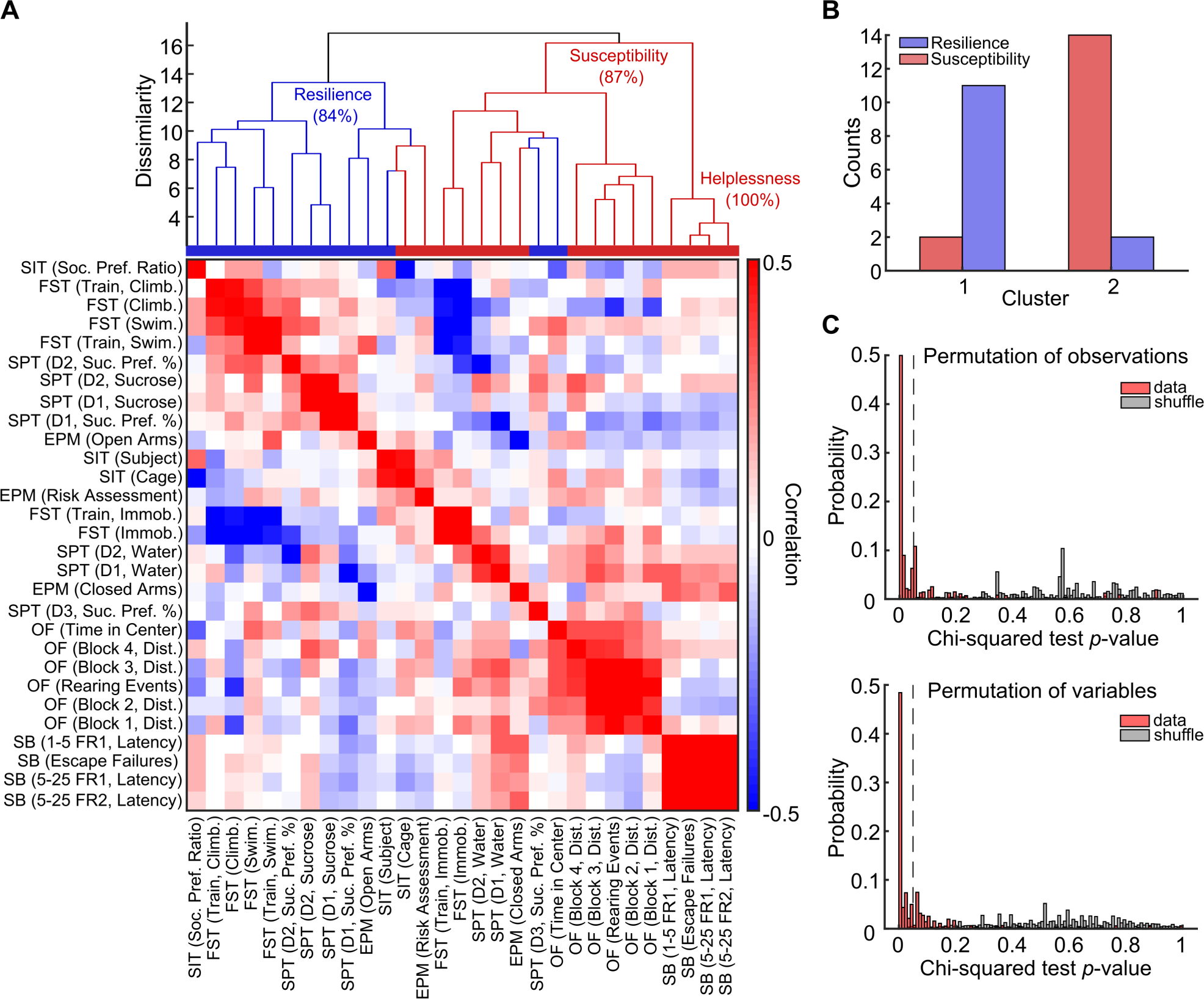
Feature clustering distinguishes behavioral variables associated with susceptibility and resilience to stress. (**A**) Hierarchical clustering of variables forms two great clusters (top) that gather behavioral attributes either associated with resilience (indicated as blue in the bar below) or susceptibility (red in the bar). The clustering is based on the correlation between the variables, which can be observed in more detail in the matrix (bottom). (**B**) The two major clusters are significantly consisted of susceptibility-(red) or resilience-related (blue) variables (X^2^_(1)_ = 15.0818, p < 0.0001, Chi-squared test). (**C**) Distributions of resilience vs. susceptibility Chi-squared test p-values across 10^4^ iterations of randomly selecting 70% of data. This permutation approach shows that the separation between resilience vs. susceptibility variables significantly occurs in most iterations even with fewer variables (top) or observations (rats; bottom) but not for shuffled data (gray bars).

Furthermore, we also observed that hierarchical clustering based on the minimum Euclidean distance separates the behavioral tests (Cophenetic coefficient.: 0.2748; X^2^_(25)_ = 101.1548, p < 0.0001, Fig. S1A) and reveals that variables are more correlated within tests than between tests (Fig. S1B). Similarly, as shown by factor analysis, seven factors suffice to capture latent covariation corresponding to the six behavioral tests, with each factor showing high loadings only across variables of the same test (Fig. S1C-D). These results demonstrate that behavioral variables are grouped into common factors representing each behavioral.

### Multidimensional patterns of covariation across behavioral measures discriminate resilience and susceptibility

Given that variables display unique patterns of organization based on covariation, our next aim was to employ Principal Component Analysis to identify which measures account for the most variance in the dataset. The analysis unveiled that the primary source of data variation is accounted for by the first three principal components, exceeding what would be expected from shuffled data (PC1: 15.01%; PC2: 14.84%; PC3: 12.21%; cumulative variance: 42.06%; see Fig. 4A). Importantly, the first PCs discriminate NH and H individuals (PC1 vs. PC2, Fig. 4B), as well as significantly distinguish NS and IS groups (PC2: t_(43)_ = −2.810, p = 0.007; PC3: t_(43)_ = 2.147, p = 0.037, Fig. 4C). Among the first PC coefficients, PC2 exhibits higher loadings of helplessness- and susceptibility-related variables, such as escape failures (SB [Escape Failures], coeff. = 0.34) and time in closed arms of EPM (EPM [Closed Arms], coeff. = 0.19), and lower loadings of resilience-related variables, such as swimming in the forced swim test (FST [Swim.], coeff. = −0.03) and sucrose preference (SPT [D1, Suc. Pref. %], coeff. = −0.33, Fig. 4D). Notably, PC1 and PC3 exhibit lower contributions to escape performance in the shuttle box (PC1: coeff. = −0.21, PC2: coeff. = −0.21) and have opposing loadings of FST (e.g., FST [immob.]: PC1, coeff. = 0.39; PC3, coeff. = 10.18), indicating independent variability in both domains. Finally, we also fitted a linear discriminant model (LDM) on PC scores. We observed better classification performance for NH vs. H compared to shuffled data, with significant improvement for PC2 (accuracy_(data)_ = 0.894, accuracy_(shuffled data)_ = 0.662, p = 0.007, LDA) and PC3 (accuracy_(data)_ = 0.907, accuracy_(shuffled data)_ = 0.656, p = 0.037, LDA, Fig. 4E). Collectively, these results demonstrate that most of the variation in the data is explained by the first components of the PCA, and these components capture the separation between NH and H individuals, as well as between susceptibility and resilience variables.

**Figure 4.**
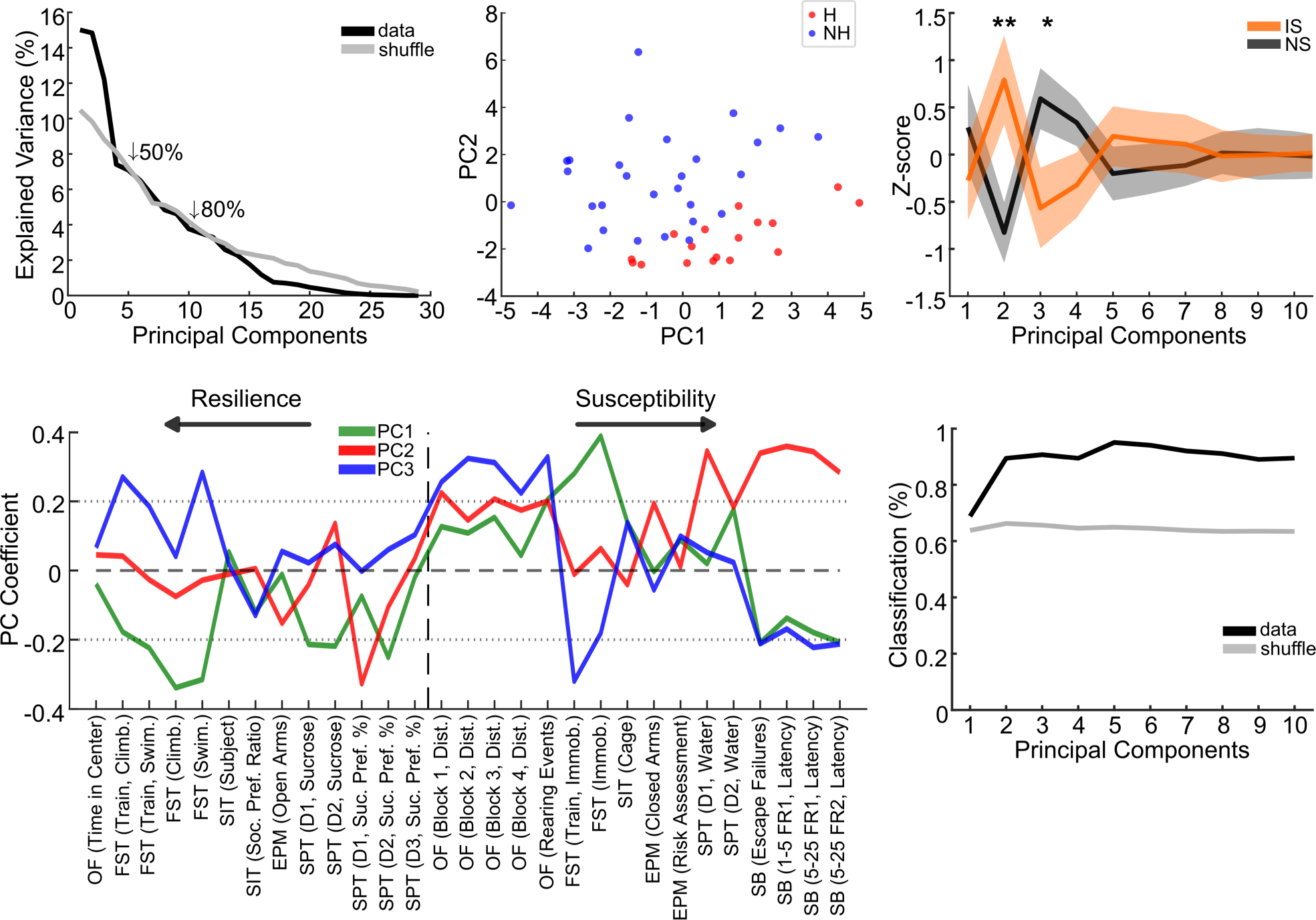
Multidimensional patterns of covariation across behavioral measures discriminate resilience and susceptibility. (**A**) Markedly greater explained variance by the first PCs, particularly PC1-3, than expected from shuffled data. (**B**) PCA map discriminates H and NH individuals. (**C**) Some of the first PCs significantly distinguish NS and IS. (**D**) Among the first PC coefficients, the PC2 exhibits high weights in helplessness- and susceptibility-related variables (right) and low on resilience-related variables (left). Also note that PC1 and PC3 exhibit lower contributions of SB escape performance and have opposing weights on FST, indicating independent variability of both domains. Variables are ordered by resilience and susceptibility, and then, by experimental design. (**E**) A linear discriminant model fitted on PC scores shows great classification performance of NH vs. H compared to shuffled data. *p<0.05, **p<0.01. Lines and shaded boundaries represent the mean ± SEM.

To assess the robustness of multidimensional covariation patterns captured by PCA, we correlated the PCs from the whole dataset with the PCs from partitioned data (70% of data randomly selected) across 10^4^ iterations. The comparison revealed that the PCs from the whole dataset also emerge in partitioned data with fewer observations (Fig. S2A), but not in the shuffled data (Fig. S2B). Additionally, by selecting the most similar PC to the whole dataset PC for every iteration, we observed a consistently high correlation of the initial PCs, both for scores and coefficients, which did not occur in the shuffled data (e.g., PC1, median values of maximum iteration correlation: Coeff. = 0.856, Scores = 0.856; Coeff._(Shuffled)_ = 0.414, Scores_(Shuffled)_ = 0.412, Fig. S2C).

Next, considering that within-test variables may be interdependent and aiming to avoid multicollinearity bias in the data, we selected representative behavioral measures from each behavioral test, opting for measures commonly used in the evaluation of neuropsychiatric disorder models (locomotion in open field; immobility in forced swim test; social interaction; time in open arms of EPM; sucrose preference; and escape failures in shuttle box). The collinearity test showed that the selected variables exhibit low interdependence (VIF values < 1.1, Fig. S3A). PCA analysis of representative variables shows that the principal source of data variation is well explained by the first three principal components (PC1: 23.82%; PC2: 21.61%; PC3: 15.78%; cumulative variance: 61.21%; Fig. S3B). PC1 is primarily described by the covariation between locomotion in the open field (OF [Total Distance], coeff. = −0.43), immobility time in forced swim test (FST [immob.], coeff. = −0.56), and sucrose preference (SPT [Average Pref.], coeff. = 0.60), while in PC2, the main coefficients of covariation are social interaction (SIT [Soc. Pref. Ratio], coeff. = 0.52) and escape failures in shuttle box (SB [Escape Failures], coeff. = 0.60, Fig. S3D). In PC3, the principal source of variation is explained by time in open arms of EPM (EPM [Open Arms], coef = 0.79) and escape failures in shuttle box (SB [Escape Failures], coeff. = 0.57, Fig. S3D). Interestingly, PC1 is sensitive to the covariation between immobility time in forced swim test and sucrose preference, two of the main measures of depressive-like behaviors. However, it is not sensitive to escape failures in shuttle box and does not discriminate animals between the groups (loading scores: IS vs. NS, t_(43)_ = −0.7881, p = 0.4349, student t-test; H vs. NH: t_(43)_ = 0.2943, p = 0.7698, student t-test, Fig. S3C). In contrast, PC2 and PC3 are described by high coefficients of escape failures in shuttle box, which covaries with social interaction in PC2 and time in open arms of EPM in PC3, measures classically associated with depressive-like behavior and anxious-like behavior, respectively (Fig. S3D). Additionally, the scores of PC2 discriminate individuals between groups regardless of clustering, with IS and H groups being better described by positive scores, indicating high social interaction and high escape failures in shuttle box (loading scores: IS vs. NS, U = 662, p = 0.0026, Wilcoxon rank-sum test; H vs. NH: U = 795, p = 0.0004, Wilcoxon rank-sum test; Fig. S3C,E). In contrast, the scores of PC3 only discriminate individuals between H and R groups, with the H group being mainly described by positive scores, indicating high time in open arms of EPM and high escape failures in EP (loading scores: IS vs. NS, t_(43)_ = 1.8122, p = 0.0769, student t-test; H vs. NH: U = 817, p < 0.0001, Wilcoxon rank-sum test; Fig. S3C,E).

### Clustering of individuals discriminates stressed subjects and reveal multidimensional behavioral phenotypes of resilience and susceptibility

PCA also revealed a wide variation in the values of PC scores associated with individuals from each group. Thus, within the same group, there are individuals whose behavioral variation is explained by more positive or negative PC scores. We performed hierarchical clustering to evaluate whether this variation is related to clusters of individuals from the general population with similar behavioral profiles. The number of clusters was determined by evaluating the silhouette value, which indicated the formation of 7 clusters (Fig. 5A). The chi-squared test revealed significant discrimination of stressed individuals by every clustering hierarchy (e.g., 7 clusters, p = 0.016, Fig. 5B) and demonstrated a significant difference in cluster composition proportions between the IS and NS groups (IS vs. NS: X^2^_(6)_ = 15.5632, p = 0.0163, Fig. 5C). The dendrogram of hierarchical clustering (HC) unveiled multivariate behavioral profiles of resilience and susceptibility (Cophenet coeff. = 0.4563, Fig. 5D). Notably, the two macroclusters displayed by the dendrogram (dissimilarity ∼= 8) exhibit a clear spectrum of distinction among individuals regarding escape failures in the shuttle box (Fig. 5E). Although most clusters present intermediate profiles of resilience or susceptibility (dissimilarity ∼= 4), both profiles are more prominently represented in Cluster 7 (susceptibility) and Cluster 2 (resilience, Fig. 5F). The susceptibility cluster stands out for displaying the most classic depressive-like behavioral alterations reported in the literature, such as reduced social interaction, reduced sucrose preference, increased immobility in the forced swim test, and increased escape failures in the shuttle box, Cluster 7, Fig. 5F). In contrast, the resilience cluster is primarily characterized by reduced escape failures in the shuttle box associated with increased sucrose preference. Additionally, there is also a cluster mainly characterized by time spent in the open arms of the EPM (Cluster 3), and a cluster in which locomotion in the open field stands out as the primary variable (Cluster 5, Fig. 5F).

**Figure 5.**
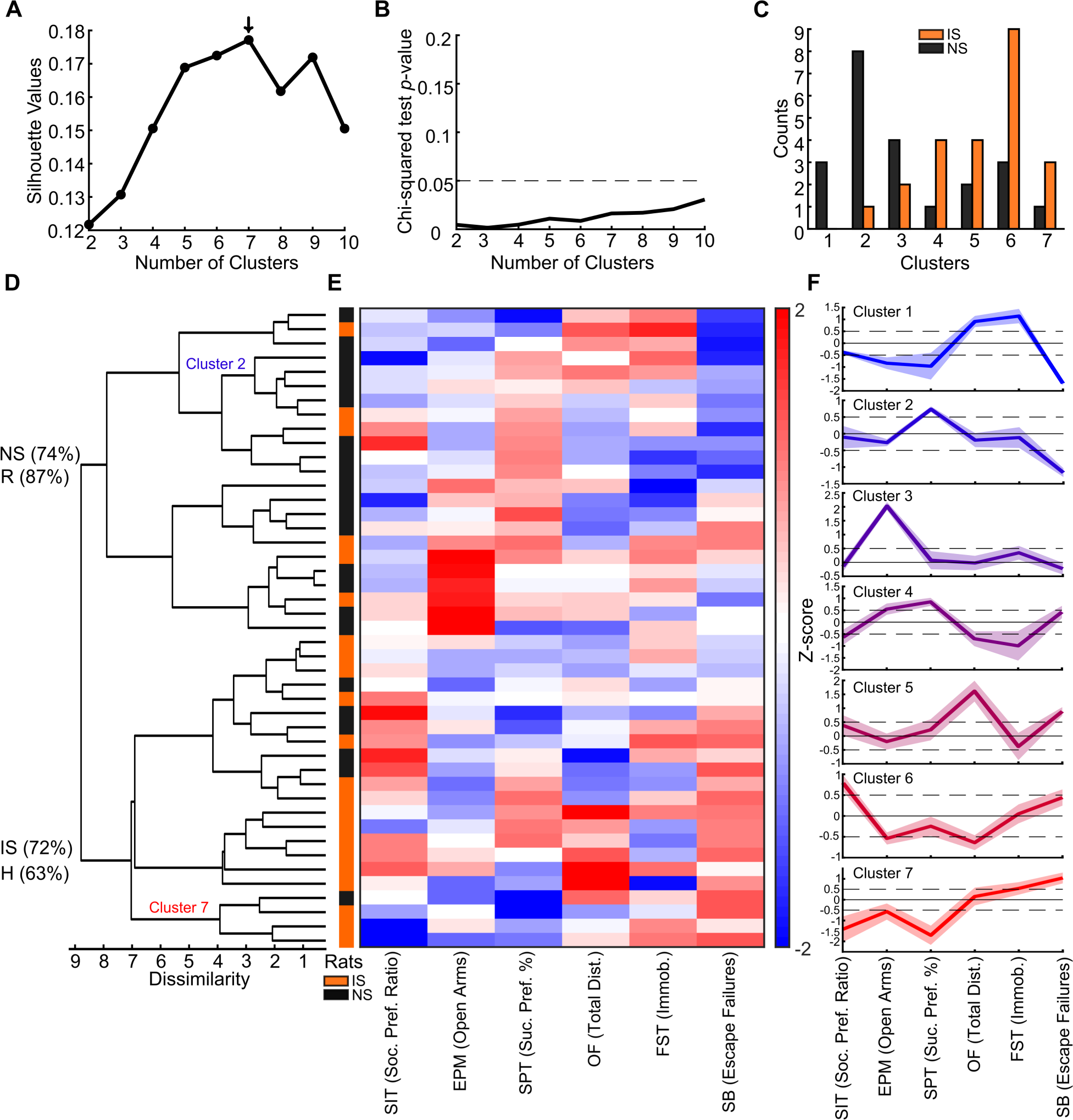
Clustering of individuals distinguish stressed individuals and reveals multidimensional behavioral phenotypes of resilience and susceptibility. (**A**) Silhouette values indicate seven clusters as optimal for hierarchical clustering. (**B**) Every clustering hierarchy significantly (IS vs. NS Chi-squared test p-value) distinguish stressed individuals. (**C**) Clusters exhibit a spectrum of distinction between proportions of NS vs. IS individuals. (**D**) Dendrogram of hierarchical clustering and (**E**) Z-scored data. Note that the two major clusters show a clear distinction at SB escape performance. (**F**) Multidimensional behavioral profiles. Note that the Cluster 7 presents a generalized profile of susceptibility. Also note that the Cluster 2 exhibits an overall profile associated with resilience. Lines and shaded boundaries represent the mean ± SEM.

To assess if different clustering algorithms reveal similar multidimensional behavioral profiles, in order to verify the robustness of the described results, we conducted unsupervised clustering analysis using also a non-hierarchical (k-means) algorithm. Silhouette values indicated that six and seven clusters were suitable for clustering (e.g., 6 clusters: Silhouette Value = 0.355, Fig. S4A), but only four to six clusters could discriminate NS vs. IS (e.g., 6 clusters: p = 0.014, Fig. S4B). Therefore, we selected six clusters as the most suitable for clustering (Fig. S4C). Remarkably, the clusters showed a spectrum of distinction in proportions between NS vs. IS individuals (X^2^ = 14.1991; p = 0.0144, Fig. S4D). Importantly, the clusters identified by k-means exhibited a high correspondence with the hierarchical clusters, notably k-means Cluster 6, identical to HC Cluster 7, both characterized by a generalized susceptibility profile (Fig. S4E). This result demonstrates that in this dataset, two different clustering algorithms reveal similar multidimensional behavioral profiles.

## Discussion

Taken together, our results indicate that behavioral variables exhibit stronger correlations within tests than between tests and organize themselves based on their relation to susceptibility or resilience to stress. This variability translates into distinct multivariate behavioral profiles, including susceptibility or resilience profiles and intermediate phenotypes, indicating multiple stress coping strategies.

Several studies have demonstrated that inescapable acute footshocks induce prolonged physiological and behavioral alterations (see Fig. S5/Supplementary Table 1). However, there is no consensus regarding the exact duration of these changes. In contrast to previous studies, our data do not confirm some frequently reported behavioral alterations that are observed over a prolonged period, such as reduced locomotion in OF (Dijken et al., 1992a; Kinn Rød et al., 2012) or time spent in the open arms of EPM (Steenbergen et al., 1991; Kinn Rød et al., 2012). On the other hand, our data are consistent with a series of studies that do not observe alterations in measures such as immobility time in FS (Ronzoni et al., 2016) or sucrose preference (Kinn Rød et al., 2012). It is essential to note that despite the similarity of the protocols (a single session of acute stress), differences in parameters, such as the number of trials and shock intensity, can yield heterogenous experimental results. These differences may also be due to the time elapsed between stress and behavioral measurement. Short & Maier (1993) demonstrated that acute inescapable footshocks (100 trials, 1 mA) lead to a reduction in social interaction in the first 48 hours, and this effect becomes non-significant three days after stress, possibly absent after seven days. This fact may explain why we did not observe alterations in the later tests of the behavioral battery, such as social preference and sucrose preference. However, it does not explain the changes observed in the startle response or in time in the open arms of EPM.

In our study, a single-day session of inescapable footshock did not elicit group-level alterations in most of the evaluated behaviors but was sufficient to induce learned helplessness and discriminate between H and NH individuals after 12 days. Interestingly, the induction of helplessness for 24 to 48 hours by acute inescapable footshock is frequently reported (Dwivedi et al., 2004; Oliveira & Hunziker, 2014; Shirayama et al., 2002; Short & Maier, 1993). However, the duration of this effect is not well-established. Some protocols report longer-lasting effects of the shock (9 days in Malberg and Duman, 2003). Notably, to our knowledge, our study is the first to show helplessness induction 12 days after acute inescapable footshock stress.

Our data reveal that reduced time in the open arms of the EPM, diminished sucrose preference, and increased escape failures, all measures captured by PC2, consistently distinguish IS from NS, as well as H from NH rats. This finding encompasses the main alterations typical of the depression-like profile reported in the literature. Indeed, the relationship between time in open arms and escape performance is widely documented. Naïve animals who spent time in open arms exhibited reduced escape performance in the shuttle box (Ho et al., 2002), indicating an association between anxiety-like phenotype and stress susceptibility. Similarly, rats selected for low sucrose consumption spend less time in the open arms of the EPM (Desousa, 1998), reinforcing the link between anhedonia and anxiety-like behavior. However, in the population of individuals in our study, the relationship between anxiety-like behavior and stress susceptibility proved to be more complex. Our results show that increased time in the open arms of the EPM and increased escape failures, both captured by PC3, also discriminate H from NH rats, indicating, together with PC2, high variability in the relationship between stress susceptibility and different levels of anxiety-like behavior.

Importantly, our results revealed that the broad inter-individual behavioral variability captured by PCA translates into clusters of individuals with multidimensional behavioral profiles. In our study, these profiles reflect different stress coping strategies, ranging from reactive to proactive coping (Koolhaas et al., 1999) and different emotional reactivity (e.g., hedonic preference). In general, some clusters are entirely represented by reactive or proactive coping strategies, while others express both strategies. This variation in coping strategies throughout the battery of tests could indicate behavioral flexibility, with the subpopulation of individuals adapting to the demands of the tests (De Boer et al., 2017). Intriguingly, both extreme clusters, the susceptibility cluster (HC Cluster 7) and the resilience cluster (HC Cluster 2), express reactive coping, despite diverging completely in escape performance in shuttle box and, consequently, in stress susceptibility. One possible explanation for this result is that while reactive coping in the susceptibility cluster may represent maladaptation (including in escape performance in shuttle box), in the resilience cluster reactive coping may represent the ability to successfully adapt to the test conditions (De Boer et al., 2017).

Interestingly, Kim et al. (2017) showed that mice subjected to a chronic protocol of inescapable shocks, regardless of susceptibility to stress, do not exhibit increased immobility time in forced swim test but rather anhedonia and anxiety-related alterations. Similarly, Stepanichev et al. (2016), using an eight-week chronic unpredictable mild stress or a two-week combined chronic stress protocol in rats, also did not observe changes in immobility time in forced swim test but instead induced anhedonia. In fact, anhedonia is one of the core symptoms of depressive disorders and is often associated with helplessness in chronic footshock stress protocols. In contrast, Meng et al. (2016) demonstrated that one-week uncontrollable foot shocks only induced learned helplessness but not anhedonia in animals. However, our data show that the relationship between anhedonia and helplessness, in conjunction with other measures, explains a portion of the data variation (PC2) and constitutes the susceptibility profile in a subset of individuals (HC Cluster 7, approximately 8.9% of the population). Indeed, HC Cluster 7 is the only cluster that exhibits the major behavioral alterations most associated with the depressive-like phenotype. Thus, it is evident that the majority of depressive-like behavioral alterations occur in a small portion of the population of individuals.

Notably, our data revealed that the immobility time in forced swim test, traditionally interpreted as a depressive-like alteration (Porsolt, 1977; Pryce et al., 2011), is not altered by inescapable footshocks. Furthermore, the immobility time shows weak correlations with other variables, and its relation with helplessness is unclear. Our data show that even individuals classified with high immobility time are unrelated to susceptibility or resilience. However, intriguingly, a subpopulation of resilient individuals (HC Cluster 1) exhibited high immobility time and a low number of escape failures, indicating that immobility may represent a possible stress-coping strategy as suggested previously (Molendijk & De Kloet, 2015, 2019; De Kloet & Molendijk, 2016). Increasingly, studies have shown that immobility does not model “despair” or helplessness, nor does it reflect depression (Molendijk & De Kloet, 2015, 2019; De Kloet & Molendijk, 2016; Commons et al., 2017). In fact, acute administration of glucocorticoids has been shown to increase immobility (Veldhuis et al., 1985), whereas the acute administration of glucocorticoid antagonists reduces immobility (De Kloet et al., 1988). Both changes are directly related to hypothalamic–pituitary– adrenal axis and not exclusively to neurobiological aspects of depression. In this sense, high immobility time in forced swim test may represent a passive strategy for energy conservation, as well as reflecting aspects of learning and memory (West, 1990). Similarly, in the sucrose preference test, the preference for water consumption over sucrose solution is interpreted as a reflex of anhedonia, one of the main symptoms of depression (Willner et al., 1987). However, the preference can sometimes be more related to other aspects of reward processing, such as motivation, than to inability to experience pleasure (Der-Avakian & Markou, 2011).

Animal models of neuropsychiatric disorders aim to replicate aspects associated with clinical symptoms rather than reproduce the entire disorder’s characteristics (Nestler & Hyman, 2010). In fact, after exposure to inescapable shock stress, only a few changes can be attributed to the development of depressive-like symptoms. Other observed alterations may have a greater significance in understanding aspects pertinent to other clinical conditions, such as anxiety and PTSD. (Christianson et al., 2010; Ronzoni et al., 2016). Therefore, applying acute inescapable footshock stress protocols should not aim to model multifaceted conditions but rather specific symptoms that may reveal shared aspects among conditions.

In the clinical setting, even patients who meet the diagnostic criteria for major depression are characterized by differences in symptom profiles and severity levels (Fried & Nesse, 2015; Maglanoc et al., 2019). This fact also points to individual biological variability as a relevant factor in the expression and relationship of depressive symptoms. Despite the majority of studies with stress-based animal models still neglecting individual variation (Einat et al., 2018), an increasing number of studies have embraced this variable and provided essential insights (Wood et al., 2010; Armario & Nadal, 2013; Ebner & Singewald, 2016; Dopfel et al., 2019; Careaga et al., 2019). In fact, our results demonstrate that inter-individual behavioral variability can reflect different stress-coping profiles. In this sense, the multivariate assessment of individual behavioral responses can reveal patterns concealed by group mean values and highlight the implications of individual variability.

The use of dimensionality reduction, clustering, and classification techniques to investigate multidimensional behavioral profiles in behavioral test batteries is a promising but still underutilized strategy (Feyissa et al., 2017; Kanari et al., 2005; Kassai et al., 2022; MacKillop et al., 2016; Rudolfová et al., 2022; Saénz et al., 2006). This approach has been applied in naive animals, allowing the characterization of inter-individual variability in different rodent strains (Feyissa et al., 2017; Kassai et al., 2022; Rudolfová et al., 2022), and in animal models of stress-related disorders (Kanari et al., 2005; Saénz et al., 2006), enabling reinterpretations of these models. Our results confirm patterns that are recurrent in these studies, such as (I) a higher correlation between measures of the same test than between tests (Feyissa et al., 2017; Kassai et al., 2022), indicating that measures of the same test may be capturing very similar alterations; (II) Each principal component (PCA) or factor (Factor Analysis) explaining only a small portion of the data variance, typically between 20% and 30% of the variance in the first component or factor (Feyissa et al., 2017; Kassai et al., 2022; Rudolfová et al., 2022), indicating that despite the existence of phenotypic subpopulations, inter-individual variability is considerably high. However, it is essential to note that in cases where the test battery measures few behavioral categories or the experimental subjects’ conditions are particular (e.g., a test battery with 3-week-old rats), the first component tends to capture a more significant portion of the data variation (Saénz et al., 2006).

Our results demonstrate that using a data-driven approach for the multidimensional assessment of behavioral profiles is a strategy that benefits from inter-individual variability and provides insights into the study of stress-related disorders. This approach has gained increasing prominence in stress research and holds a significant position in the future of rodent models in depression research (Gururajan et al., 2019).

## Conclusions

We have demonstrated that the multivariate assessment of behavioral responses to acute inescapable footshocks can reveal susceptibility or resilience behavioral profiles and intermediate phenotypes. Thus, our approach emphasizes the importance of considering individual variability and the relationship between multiple behavioral measures to understand different behavioral responses to stressors, providing new insights into studying animal models of neuropsychiatric and stress related disorders.

## Methods

### Animals

We utilized forty-eight male Wistar Hannover rats, aged seven to ten weeks old (weight range: 250g - 300g), obtained from the Central Animal Facility at the University of São Paulo. Rats were housed under standard conditions in the animal facility, with four rats per cage based on their respective experimental groups. The temperature was maintained at 24 ± 2 °C, and the rats had *ad libitum* access to food and water. A 12-hour light/12-hour dark cycle was implemented, with lights on at 7:00 a.m. Handling was carried out two days before the experimentation.

### Experimental Design

Rats were randomly divided into a ‘not shock’ group (NS; not exposed to stress before the test battery; n = 24) and an ‘inescapable shock’ group (IS; exposed to inescapable electric footshock before the test battery; n = 24). Subsequently, they were subjected to a comprehensive behavioral battery test on consecutive days. The habituation to the experimental room was carried out previously to each test and lasted for 20 minutes. Lighting in the rooms was kept constant during all procedures. All apparatus were cleaned with an alcohol solution (variable concentration, between 30% to 70%, depending on the test performed) and deionized water. Experiments were carried out in the light cycle, 12:00 a.m. to 6:00 p.m., except for inescapable shock sessions starting at 7:00 a.m. and ending at 6:00 p.m. due to the duration of each session (1 hour and 20 minutes per animal). All behavioral tests were video recorded, and the quantification was performed using the X-Plo-Rat 2005 1.1.0 software (Tejada et al., 2018). The analysis was conducted blindly, without knowledge of the experimental groups. All procedures were approved by the local ethics committee on animal experimentation (Ribeirão Preto Medical School, University of São Paulo; protocol number: 78/2020.

### Behavioral test battery

The test battery was composed of tests classically used in behavioral neurosciences to investigate the effects of stress.. Tests were performed on consecutive days, according to the following sequential order: open field, forced swim, social preference, elevated plus maze, object recognition, prepulse inhibition/startle, sucrose preference, and escape performance. The most potentially stressful tests were performed at the beginning of the test battery to reduce possible confounding effects on the escape performance.

### Inescapable footshocks

Rats were exposed to inescapable footshock in the active avoidance shuttle box adapted apparatus (Insight). The apparatus consists of a floor with metal bars, LEDs for emitting light stimulus, infrared light emitters coupled to animal location detectors, and a shock generator with an electric current limiter. The apparatus has dimensions of 30.7 cm in height x 33 cm in width x 54 cm in length and was placed in a sound-attenuating chamber acoustic insulation box (51 cm in height x 48 cm in width x 63 cm in diameter).

The IS group was exposed to 60 trials of electric foot shocks. Each trial started with exposure to a luminous stimulus (200 lux; conditioned stimulus) for 20 seconds. Ten seconds after the stimulus started, the animals received inescapable shocks with intensity of 0.8 mA and uninterrupted duration of 10 seconds. Both stimuli ended simultaneously and were followed by a 40 ± 20 seconds interval between trials. The NS group was submitted to the same protocol but received no shocks. The entire experiment was carried out in a room without lighting (0 lux).

### Open Field Test (OF)

The OF is used to assess locomotor and anxiety-like behavior of animals when exposed to an open environment (Seibenhener & Wooten, 2015). The apparatus consists of an acrylic floor arena (Insight, 46 cm in height and 46 cm in length) surrounded by glass walls. In this test, the animals were placed individually in the arena center to explore the environment freely for 20 minutes. Throughout the task, the lighting intensity in the room was kept constant (500 lux). The movements were captured by infrared sensors detecting presence and movement arranged on the walls of the arena.

### Forced Swim Test (FST)

The FS is traditionally used to assess behaviors associated with the depressive-like phenotype. Longer immobility time and shorter escape-oriented behaviors are measures associated with the depressive-like phenotype (Slattery & Cryan, 2012). In this study, we carried out the test in two stages, each in a day, under the same environmental conditions (constant lighting of 500 lux). On the first day, the animals were placed in the center of a transparent acrylic cylinder (60 cm in height and 30 cm in length, filled with water up to 30 cm in height at 22 ° C) for 15 minutes. Twenty-four hours later, the animals were again subjected to the task in the same settings for 5 minutes. After each session, the rats were carefully dried before being returned to their cages. The immobility time, swimming time, escape time, immobility latency, diving events, and fecal cakes production were considered for analysis.

### Social Interaction Test (SIT)

The SIT evaluates sociability behaviors represented by the approximation or avoidance between the animals tested (Berton et al., 2006). In our study, the test was adapted from Lukas et al. (2011) and carried out previously described open field arena. Rats were initially habituated to the apparatus for 30 seconds and then to the presence of an empty cage (12.5 cm in height and 9 cm in width) positioned against a wall of the arena for 5 minutes. Subsequently, the tested rat was removed from the arena for 2 minutes to accommodate the co-specific animal (same strain, sex, age, size and weight) in the cage. The tested animal was reinserted in the arena for 10 minutes, and all of its movements were filmed with an infrared camera. The experiment was carried out under constant red lighting, not visible to rats, with other lights turned off (7 lux). We considered for analysis the time spent in interaction with the empty cage or the cage filled by the co-specific animal, and the production of fecal boli. The interaction behavior was determined as the interval time in which the tested animal head was directed to the cage, and its muzzle was at a distance of approximately 1 cm or less.

### Elevated Plus Maze Test (EPM)

The elevated plus maze test is used to assess behaviors associated with the anxious-like phenotype (Walf & Frye, 2007). The test generates in the animal the conflict between environmental novelty exploration and innate fear of open places. More time spent in closed arms and avoiding open arms are associated with the anxious-like phenotype. The behavioral apparatus (80 cm in height, 110 cm in width and 110 cm in length) consists of four raised arms of the same size intersecting in parallel at a central point. Two arms are closed by side walls and the other two are open. The apparatus is 30 cm away from the room walls, in an environment without visual, olfactory or auditory clues and under constant exposure to homogeneous lighting (2 lux). In this test, animals were placed over the center of the apparatus and filmed with an infrared camera for 5 minutes. For analysis, we consider the period of stay and entry events in each arm and in the center, as well as risk assessment, rearing, vertical exploration, head dipping, and fecal cake production behaviors.

### Novel Object Recognition Test (NOR)

The NOR assesses the recognition memory based on the exposure of a familiar object maintained throughout the sessions and a new object that varies with each session. A low discrimination ratio of the new object compared to the familiar object is associated with cognitive deficits (Bevins & Besheer, 2006). In this study, the task was performed in a closed arena (83 cm in height x 42.5 cm in width x 39.5 cm in length), with its lighting (150 lux) and an air exchange system (fan attached to the arena wall) over three days. On the first day, animals were habituated to the apparatus for 20 minutes. Twenty-four hours later, the habituation lasted 10 minutes and was followed by exposure to two identical objects (objects A and A’) inserted in the arena for 5 minutes. After 15 minutes interval outside the arena, the animal was reinserted in the apparatus for 5 minutes and exposed to objects (A, familiar object, and B, new object). In this stage, we tested the animal’s short-term memory based on the discrimination between objects A and B. The next day, the reexposure to objects (A, familiar object, and C, new object) lasted 5 minutes. At this stage, we tested the animal’s long-term memory and its discrimination between familiar and new objects. The entire experiment was carried out under constant lighting and was filmed. For analysis, we considered the relative and the total time of object interaction. We calculated the new object discrimination index according to the formula: TN/(TF + TN), where TN = new object exploration time and TF = familiar object exploration time.

### Prepulse Inhibition and Acoustic Startle Reflex Test (PPI/Startle)

The PPI/Startle assesses the startle motor response caused by an intense sound stimulus (pulse). This response is reduced when the pulse is preceded by a less intense sound stimulus, the prepulse. (Fendt & Koch, 2013). In this test, the animals were exposed to pulses, prepulses, prepulses + pulses, and only evaluated for their startle response or inhibition. The sound stimuli were generated by two loudspeakers located inside an acoustic isolation box. The animals were restricted to a cage (12.5 cm in height, 9 cm in width and 19 cm in length) coupled to a stabilimeter that measures the motor reaction. After the initial 5 minutes of habituation to the apparatus, the animals were subjected to 60 semi-randomized sound stimulus trials between pulse stimuli (6 stimuli; 120 dB intensity; 40 ms duration), 18 prepulse stimuli (71, 77 and 83 dB intensity; 6 stimuli for each intensity; 20 ms duration), 18 prepulse + pulse stimuli (pre-pulse at 71, 77 and 83 dB intensity; 6 stimuli for each intensity; 20 ms duration, followed after 100 ms by the pulse of 120 dB of intensity and 40 ms of duration) and only background noise stimulus (6 trials; 65 dB of intensity). The interval between each trial lasted 15 ± 8 seconds (adapted from Ruggiero, 2016). The lighting of the room (500 lux) and from inside apparatus (0 lux) remained constant during the experiment. The following behavioral measures recorded by the software integrated into the apparatus (Insight) were considered for analysis: the pulse trials mean (represented as startle reflex amplitude), the startle habituation (HAB) percentage (%HAB=100-(100*(HAB1/HAB0))), and the prepulse inhibition percentage for three stimulus intensities (%PPI=100-(((PP+P)/P)*100)).

### Sucrose Preference Test (SPT)

The SPT is classically used as an indicator of anhedonia, the main symptom of the depressive phenotype. The absence of preference for sucrose compared to water is associated with anhedonia (Liu et al., 2018). In this test, animals were housed individually in homecages for 72 hours with *ad libitum* access to two drinking fountains: one containing only water and other containing 1% concentrated sucrose solution, both filled with 500 ml. Every 24 hours, the volume of liquid remaining was measured, and the drinking fountains were immediately switched positions (left vs. right) in the homecages. The animals were handled only for weighing, and there was no additional exposure to lighting.

### Shuttle Box (SB)

The SB with escapable shock evaluates the escape response to controllable stress. In this test, the shock duration is controllable, in contrast to the inescapable shock whose duration is fixed (Maier & Seligman, 2016). For the shock termination, animals must jump over a 1 cm height bar obstacle (an escape response) placed in the shuttle box center or wait for the 10 second period to end. The high number of escape failures as well as high mean escape latencies are interpreted as learned helplessness indicators. In our study, the task consisted of 30 escapable shock trials with 0.6 mA intensity. Each trial consisted of a light stimulus lasting 10 seconds, followed by a light stimulus + shock lasting 10 seconds and, finally, an interval of 40 ± 20 seconds without application of stimuli. The 5 first trials had a fixed ratio-1 (FR-1) task acquisition criterion, in which shock termination was conditioned to the execution of a single escape response performed during the shock duration interval (10 s). The subsequent 25 trials had a fixed ratio-2 (FR-2) in which the termination of shock was conditioned to two escape responses. The experimental room remained without lighting (0 lux), except for the sporadic luminosity of the apparatus’ light stimulus (200 lux), and animal movements were filmed with infrared camera. For analysis we considered FR-1 mean escape latencies, FR-2 mean escape latencies and the number of escape failures, all the measures obtained by the software integrated into the apparatus (Insight). Escape latencies are defined as the time interval between the shock onset and each escape performed.

### Statistical Analysis

We used the Kolmogorov test to infer a normal distribution between experimental groups. We used the t-student test to compare samples following a normal distribution, and we applied the Wilcoxon rank-sum test as a non-parametric alternative. These procedures were performed using custom MATLAB (R2022b, Mathworks) code. Data are represented as the mean ± standard error of the mean (SEM) for line and bar plots. Box-plot data are represented as 1st quartiles, medians, and 3rd quartiles, with whiskers representing the extreme values.

### Multivariate Analysis

Our multivariate analysis primarily focused on investigating the relationships between data variables across different behavioral tests. The aim was to uncover patterns that could reveal distinct phenotypes characterized by specific behavioral traits or factors that might not be evident when examining population-level averages alone. To determine the most relevant behavioral variables for each test, we considered theoretical principles, incorporating classical variables from the literature, and encompassing various behavioral categories, such as anxiety measures, recognition memory, sociability, anhedonia, sensorimotor filter, helplessness, and exploitation.

All multivariate analysis was performed in MATLAB using custom codes and built-in algorithms. Initially, all variables of each animal were normalized by subtracting the mean of the samples from both groups and dividing by the standard deviation of the mean (z-score) for each behavioral variable.

### Linear Correlation

We used Spearman’s correlation to identify linear dependence between the variables, obtaining the p-value and the correlation coefficient (r_s_). In the construction of correlation matrices, the color scale represents the values obtained from the coefficients. To facilitate visualization, the diagonal axis of the matrices was defined as zero, representing the correlation of each variable with itself.

### Principal Component Analysis (PCA)

PCA is a dimensionality reduction method that summarizes multivariate data into a smaller set of variables derived from the original set. The original data multidimensional projections (in our study different behavioral variables) are rotated to obtain new axes containing the largest variance in the data set. The axes are orthogonal to each other, thus the covariance between the axes is zero (the main axes are not correlated). In this sense, the new axes projections (main components) can help reveal variance patterns in multidimensional data. In our data, we perform the PCA using the singular value decomposition of the MATLAB PCA function. After the identification of the main components (PCs), the data were projected on new axes where the coefficients of the linear combination (loadings) of the original variables from which the principal components (PCs) are constructed were calculated. The PC loadings were compared between the conditions (IS vs. NS and H vs. R groups).

### Clustering Analysis

Clustering analysis is used to group variables (e.g. behaviors) or observations (e.g. individuals) without previously defining classes (without supervision). These algorithms try to find similar data and merge them into a single cluster. In this way, data presenting more similar characteristics are grouped into a class. In our study, we employed k-means clustering on the observations (individuals) to assess the formation of multivariate behavioral profiles, and we utilized hierarchical clustering on the variables (behaviors) to evaluate the relationship between behavioral measures. The k-means clustering method separates data into a pre-defined number of clusters using a distance metric between the data and centroids, with centroids initially established randomly or pre-defined, and iteratively recalculated by taking the mean of associated points until convergence is achieved. In our study, we used the quadratic Euclidean distance as a measure, as well as random values of the initial centroid. The choice of the cluster number was based on silhouette values (a measure of the proximity to other data in the cluster in relation to the data closest to a neighboring cluster). We repeated the clustering method a thousand times, since as the initial centroid values are random, the convergence can occur in local minimums, and we choose the result that presented the lowest intra-cluster variance. In addition, hierarchical clustering algorithms were employed, which establish a hierarchical relationship among the analyzed data by connecting closer data pairs based on a distance measurement. These clusters are subsequently merged into larger clusters, and the distances between clusters and all other points are calculated, with the closest cluster pairs continuously connected. The linear correlation coefficient between the cophenetic distances derived from the hierarchical tree and the original distances (or dissimilarities) used to construct the tree determines the cophenetic correlation, which characterizes the hierarchical relationships among clusters. In our study, we used Spearman’s correlation values between behavioral variables as a distance measure and Ward’s method of minimum variance to establish the link between clusters.

## Data availability

The data are available from the corresponding author on reasonable request.

## Acknowledgements

This research was funded by Fundação de Amparo à Pesquisa do Estado de São Paulo - FAPESP (DBM: 2022/16812-3; MTR: 2020/01510-6; RNR.: 2018/02303-4; JPL: 2016/17882-4), Coordenação de Aperfeiçoamento de Pessoal de Nível Superior - CAPES - Finance Code 001 (BAOJ: 88887.469138/2019-00 and 88887.820541/2023-00; DBM: 88882.328283/2019-01), and Conselho Nacional de Desenvolvimento Científico e Tecnológico - CNPq (TP: 164165/2018-5; JPL: 305104/2020-9 and 422911/2021-6). We thank Antonio Renato Meirelles e Silva for technical support.

## Author contributions

RNR conceived the research and supervised the project; BAOJ, DBM, MTR, TP and RNR designed the methodology; BAOJ, DBM, and RNR performed the formal analysis and prepared the figures; MTR and TP contributed to analysis and data curation; BAOJ wrote the original draft and drawn the illustrations; JPL provided resources and financial support. All authors revised, edited, and approved the manuscript.

## Supplementary Material

### Supplementary Figures and Legends

**Figure S1.**
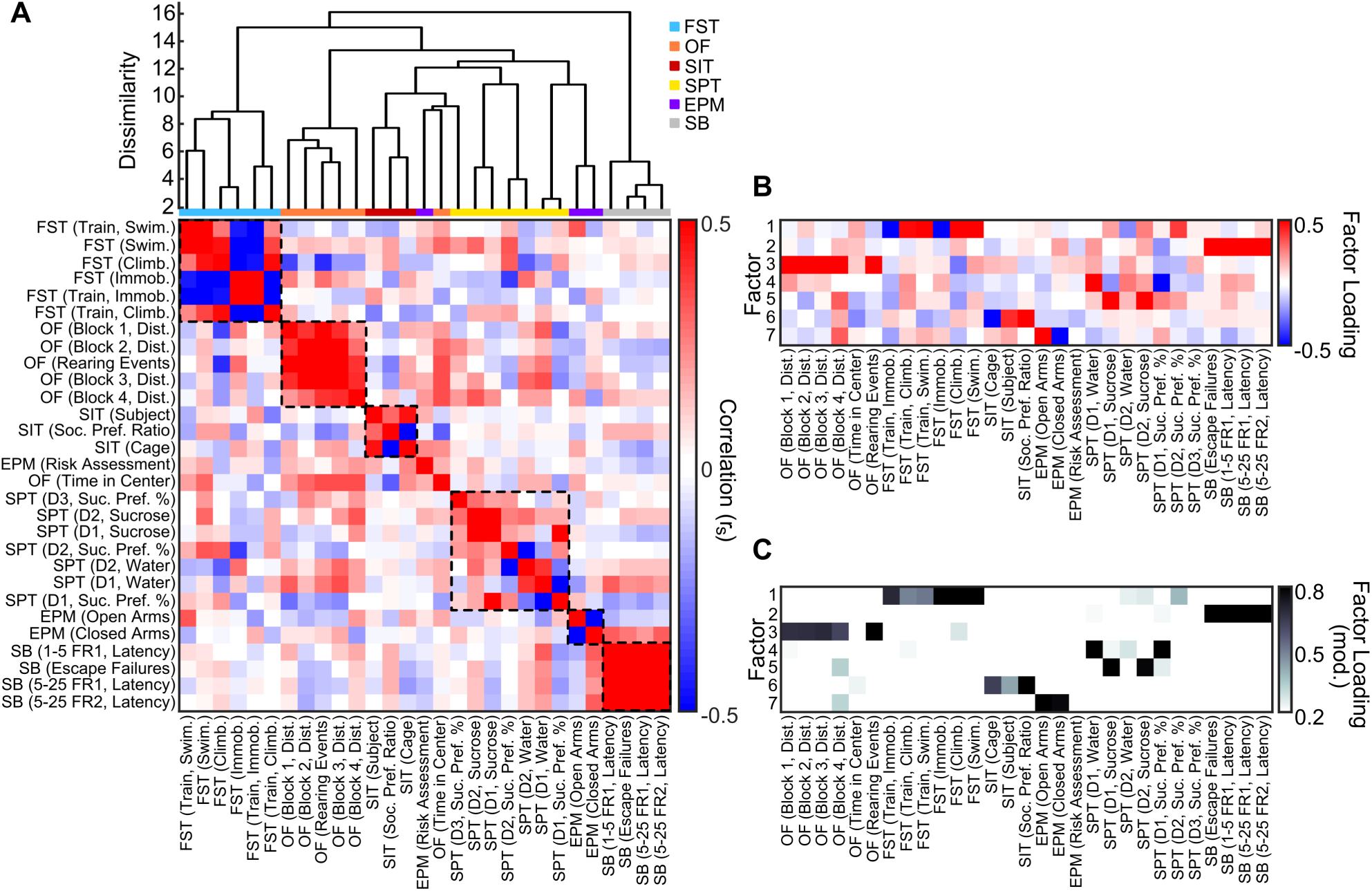
Stronger intra-test than inter-test behavioral correlations. (**A**) Feature clustering by sign-independent pattern similarity (minimum Euclidean distance) clearly separates the behavioral tests (each test is indicated by a color in the bar below). Dashed lines in the correlation matrix indicate clustered variables of the same test. (**C-D**) Factor analysis for seven factors effectively captures latent covariation corresponding to the six behavioral tests. Each factor shows high loadings only across variables of the same test. Variables are ordered by the experimental design.

**Figure S2.**
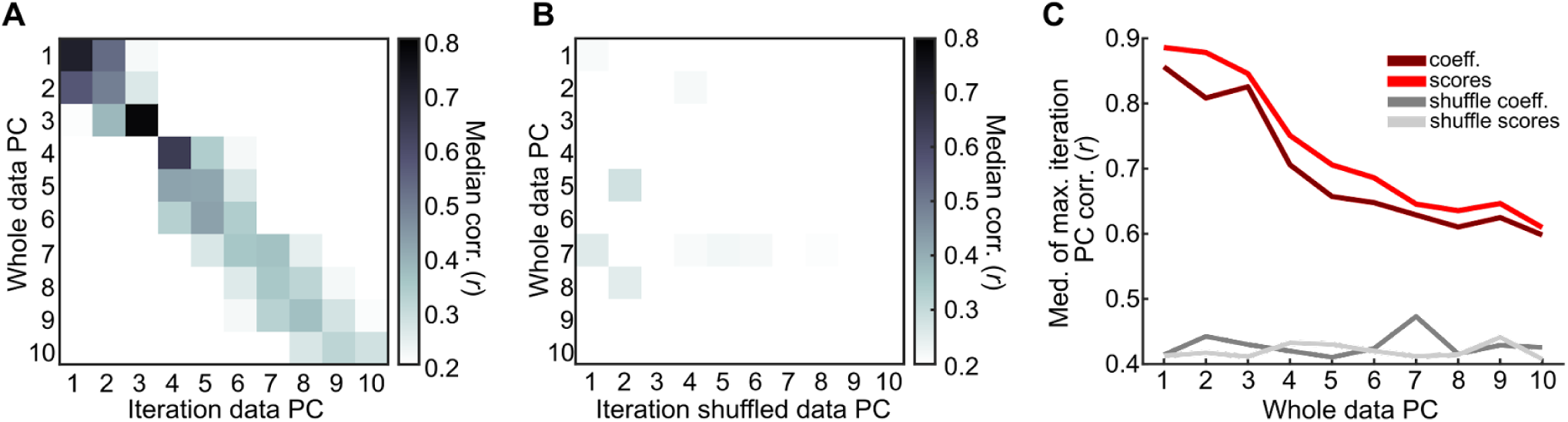
Robustness of multidimensional patterns of covariation. (**A**) The PCs from the whole data also emerge in partitioned data with fewer observations. The heatmap indicates that the PC’s coefficients are frequently correlated to PCs of partitioned data (randomly selected 70% of observations across 10^4^ iterations). Note that each PC, but especially the first ones, shows great correlation with PCs of close ranking of variance explained. (**B**) There are no clear correlations between whole data PCs to that from shuffled data. (**C**) By choosing the most similar PC to the whole data PC for every iteration, we observed a frequent high correlation of the initial PC’s, for both scores and coefficients, that does not emerge in the shuffled data.

**Figure S3.**
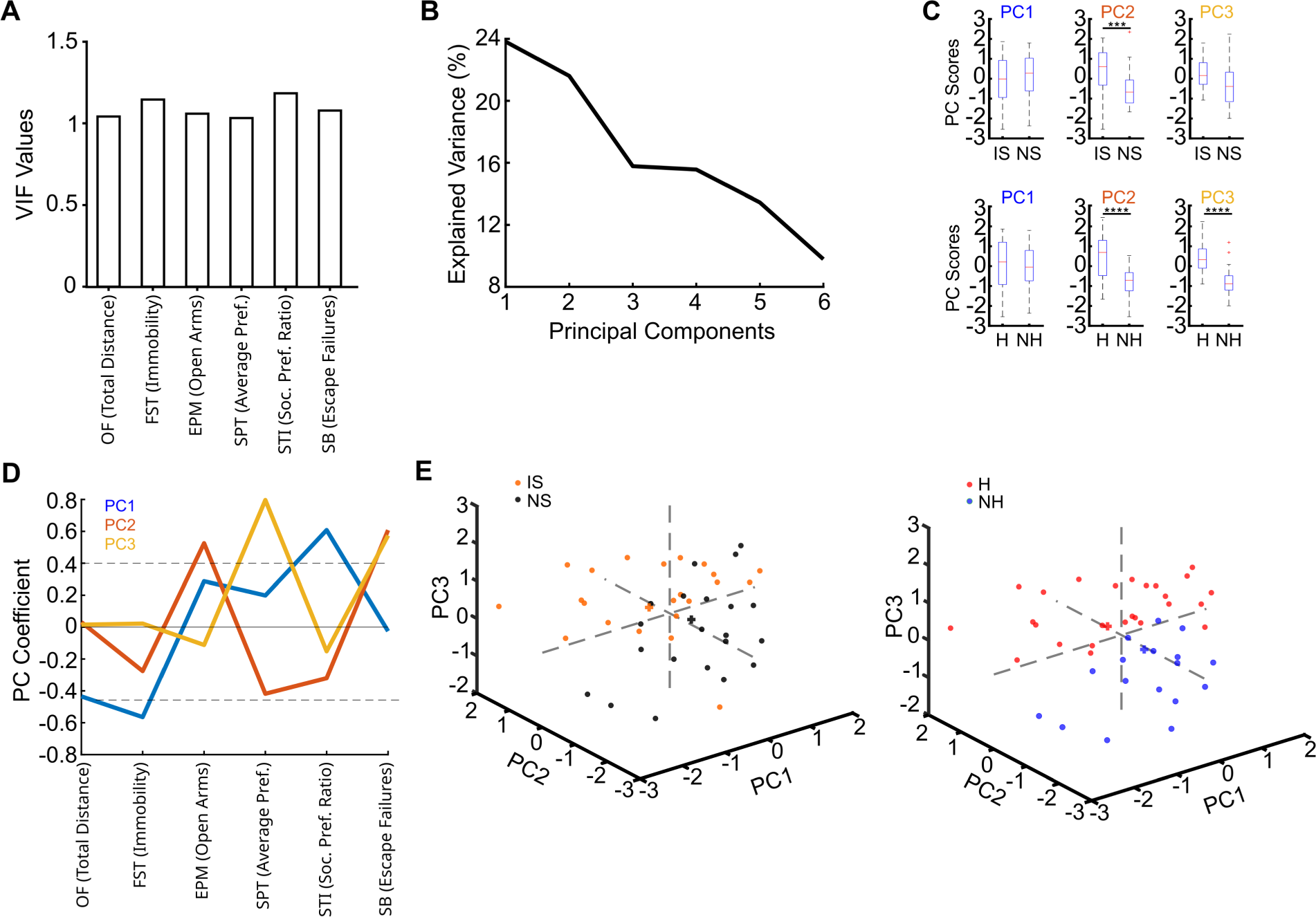
Reduced time in the open arms of the EPM, diminished sucrose preference, and increased escape failures, consistently distinguish susceptible from resilient rats. **(A)** Collinearity evaluation. Variance inflation factor (VIF) values indicate low interdependence between the selected variables. (**B**) Explained variance of the Principal Components. (**C**) Scores of individuals by group on each Principal Component. The groups are distinguished by PC2 (IS vs. NS, H vs. NH) and PC3 (H vs. NH). Student t-test or Wilcoxon rank-sum test, ***p < 0.001, ****p < 0.0001. (**D**) Coefficients of principal components related to behavioral variables. **(E**) The scores of individuals were projected onto the axes space of the first three Principal Components. Individuals categorized between IS and NS groups (left) or H and NH clusters (right).

**Figure S4.**
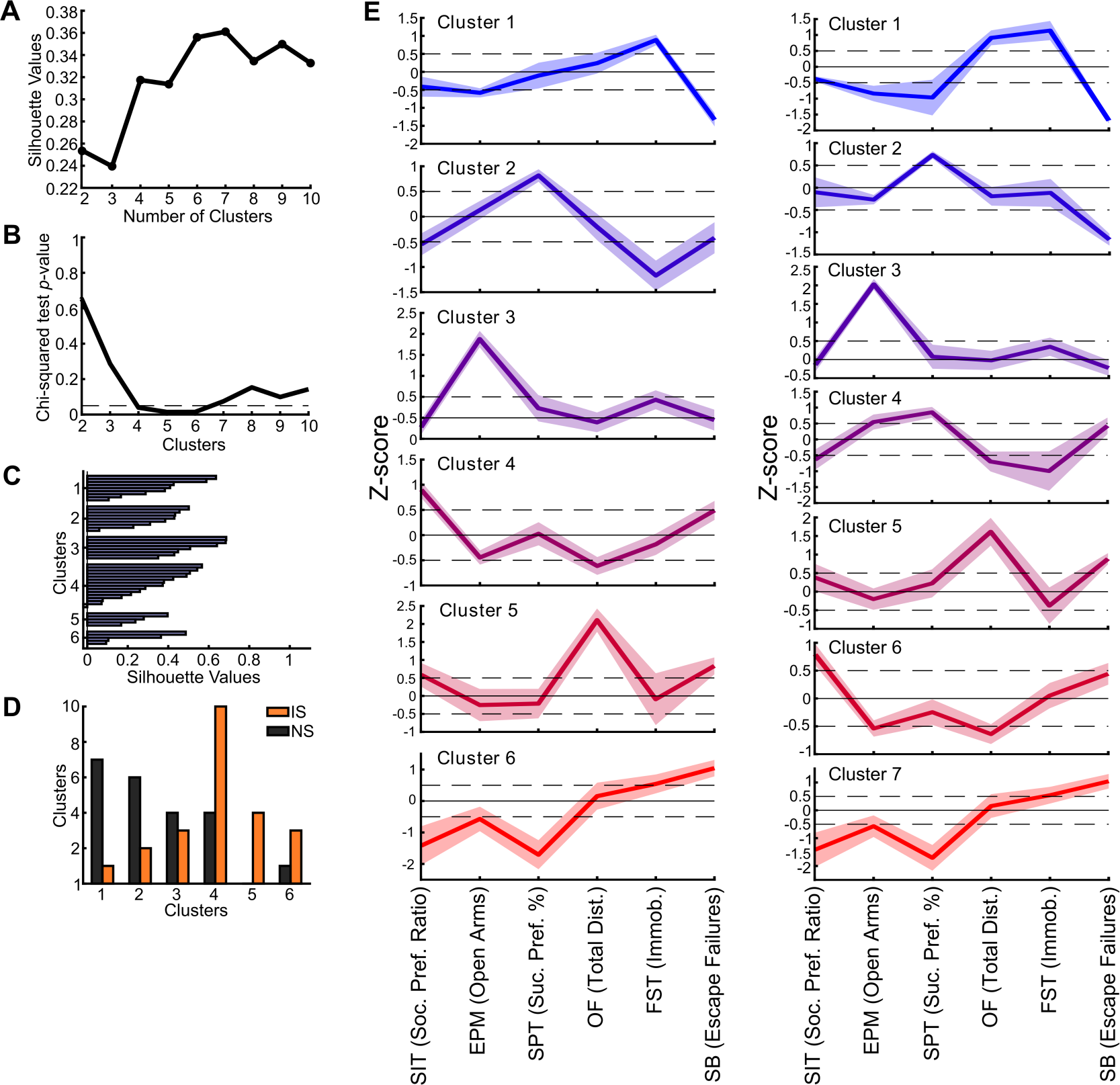
Distinct clustering algorithms reveal similar behavioral profiles. (**A**) Silhouette values indicate six and seven clusters as appropriate for k-means clustering. (**B**) Specifically, four to six clusters can discriminate between NS vs. IS (IS vs. NS Chi-squared test p-value). (**C**) Silhouette values for six clusters. (**D**) Clusters show a spectrum of distinction between proportions of NS vs. IS individuals. (**E**) Multidimensional behavioral profiles identified by k-means (six clusters, left) and hierarchical (seven clusters, right) clustering. Note the correspondence (indicated by similar color) of the multidimensional profiles between the two distinct clustering algorithms. Also note the emergence of an identical grouping as the HC Cluster 7 characterized by a generalized susceptibility profile. The HC clusters are the same as in Figure 5. Variables are ordered by resilience and susceptibility, then by experimental design. Boundedline represents the mean ± SEM.

**Figure S5.**
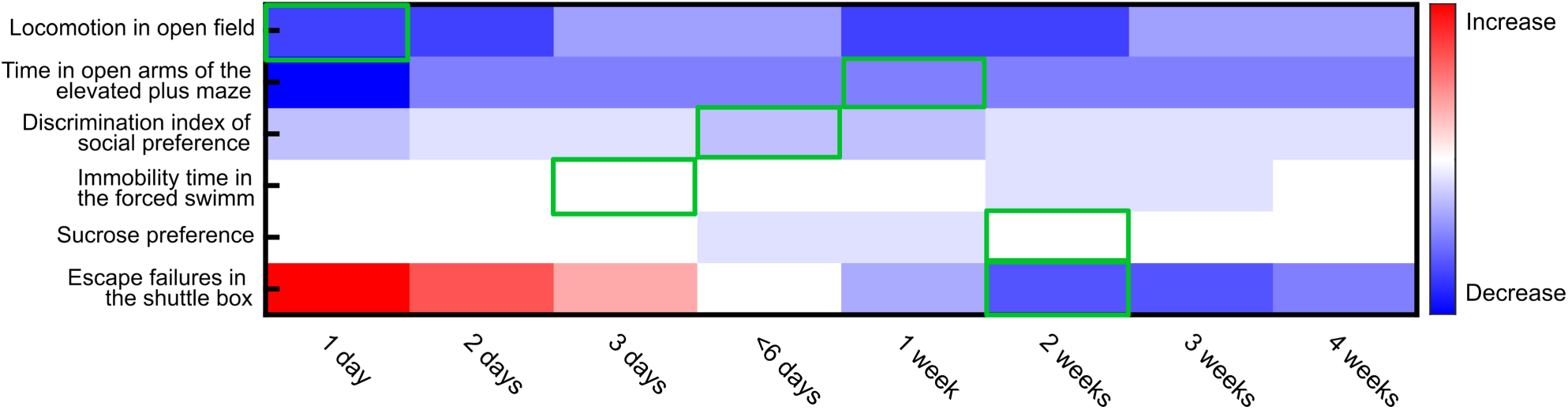
Literature reported behavioral effects of acute inescapable shocks. All comparisons are against the respective control groups. Green boxes indicate when the respective behavioral tests in our study were conducted.

**Supplementary Table 1.** Table containing the data and references for Figure S5.

| Behavioral test | 1 Day | 2 Days | 3 Days | < 6 Days | 1 Week | 2 Weeks | 3 Weeks | 4 Weeks | Authors |
| --- | --- | --- | --- | --- | --- | --- | --- | --- | --- |
| OF | ↓ | ↓ |  | ↓ | ↓ | ↓ |  |  | Dijken et al. (1992a) |
| OF |  |  |  |  |  |  |  | ↓ | Dijken et al. (1992b) |
| OF |  |  |  |  | ↓ | ↓ |  |  | Kinn Rød et al. (2012) |
| FST |  |  |  |  |  | ↓ |  |  | Dijken et al. (1992b) |
| SIT | ↓ |  |  |  |  |  |  |  | Christianson et al. (2010) |
| EPM |  |  |  |  | ↓ | ↓ | ↓ |  | Rod et al. (2012) |
| EPM |  |  |  |  |  | ↑ |  |  | Belda et al. (2004) |
| EPM | ↓ |  |  |  |  |  |  |  | Steenbergen et al. (1990) |
| EPM | ↓ |  | ↓ |  |  |  |  |  | Steenbergen et al. (1991) |
| SPT |  |  |  | ↓ |  |  |  |  | Kinn Rød et al. (2012) |
| SB | ↑ |  |  |  |  |  |  |  | Shirayama et al. (2002) |
| SB |  |  |  |  |  | ↓ | ↓ |  | Dijken et al. (1992b) |
| SB |  | ↑ |  |  |  |  |  |  | Dwivedi et al. (2004) |
| SB |  | ↑ |  |  |  |  |  |  | Dwivedi et al. (2005) |

